# Cellular and molecular networks of intercellular signaling in bone marrow hematopoiesis

**DOI:** 10.1101/2024.08.25.609544

**Authors:** Zachary V. Thomas, Bowen Wang, Wade R. Boohar, Mary Vergel, Jiya Eerdeng, Dayeon J. Shon, Michael B. Elowitz, Rong Lu

## Abstract

Hematopoietic stem and progenitor cells (HSPCs) rely on intercellular signaling to maintain and adjust their production of blood and immune cells. This process occurs in the semi-fluidic bone marrow, hosting dozens of cell types that constantly migrate and interact. To elucidate the dynamic network of cell-cell interaction and signaling transduction underlying hematopoiesis, we developed an algorithm to measure Cell-cell Spatial Interaction Probability (CellSIP) by integrating data on ligand and receptor expression, cell type abundance, and cellular spatial positioning. Using new and published mouse datasets, we validated CellSIP and uncovered signaling transductions indicating feedback mechanisms underlying hematopoiesis. Moreover, we identified significant correlations between signaling pathways across individual HSPCs at the same hematopoiesis stage. These pathway correlations illuminate the organization of cellular and signaling networks underlying hematopoiesis, revealing new regulators through their associations with established ones. The signaling quantification and correlation data are available through the Hematopoiesis Intercellular Signaling Explorer (HISE).

## INTRODUCTION

In vertebrates such as mice and humans, blood and immune cell regeneration is carried out by hematopoietic stem and progenitor cells (HSPCs) within the bone marrow.^1,2^ HSPCs rely on intercellular signaling to maintain homeostatic hematopoiesis and adjust their production of blood and immune cells in response to injury and disease.^3,4^ Previous studies have uncovered several signaling pathways that modulate HSPCs, including the Kit, Midkine (MK), Collagen, and Adiponectin pathways.^5–9^ Additionally, research has revealed key bone marrow cell types, such as mesenchymal cells and endothelial cells, which interact with and regulate HSPCs.^10–14^ Emerging evidence suggests that HSPCs not only passively receive, but also actively send signals.^15,16^ However, past research has primarily focused on individual signaling pathways, leaving a poor understanding of the overall signaling environment, including the crosstalk between pathways and the relative contributions of each signal. This knowledge gap is critical, because the bone marrow contains over three dozen cell types that constantly migrate and interact with each other. HSPCs continuously receive and send numerous signals that collectively determine their cell fates.^17,18^ Some signaling pathways may crosstalk with one another. Combinatorial signals have been shown to dictate distinct regulatory responses that are not simply the sum of individual signaling pathways.^19–22^ Therefore, applying a quantitative systems biology approach to unravel the intercellular signaling network of HSPCs holds promise for uncovering novel regulators and elucidating key regulatory mechanisms.

Profiling and quantifying the intercellular signaling of HSPCs is challenging. Unlike other tissues, the bone marrow is semi-fluidic without a fixed structure.^23–26^ Tracking the dynamic interactions of dozens of cell types is difficult due to the large number of biomarkers required for annotating cell identities. In addition, HSPCs constantly mobilize in the bone marrow, as well as in and out of peripheral blood circulation.^24,27–29^ Their mobilization is not random. For example, hematopoietic stem cells (HSCs) are frequently found in specific types of bone marrow microenvironments such as the endosteal-perivascular niche, which is mediated by chemokine-receptor interactions such as the Cxcl12/Cxcr4 pathway, as well as cell adhesion signaling such as the Vcam1/VLA-4 pathway.^14,17,30–32^ The spatial positioning of HSPCs within the heterogeneous bone marrow environment dictates their exposure to distinct signaling molecules. Moreover, heterogeneous gene expression of individual HSPCs^33^ results in varying sensitivity to intercellular signals and differences in the signals they send to other cells.

Taking into account the heterogeneous gene expression of HSPCs, their constant migration, and the diverse bone marrow environment, here we have developed an algorithm to measure Cell-cell Spatial Interaction Probability (CellSIP) in mouse bone marrow. Using CellSIP, we have systematically identified and quantified signaling interactions between HSPCs and major cell types in the mouse bone marrow. We have also uncovered significant correlations between these intercellular signals across individual HSPCs. These analyses provide a comprehensive genome-wide profile of HSPC signaling interactions, which can be accessed through the Hematopoiesis Intercellular Signaling Explorer (HISE).

## RESULTS

### CellSIP quantifies cell-cell signaling by integrating spatial interaction analysis

To identify and quantify the intercellular signaling underlying hematopoiesis in the bone marrow, we have developed an algorithm called CellSIP by integrating multiple data sources including molecular interaction intensity, cell type frequency, and cellular spatial positioning (**Figure 1A**). Specifically, CellSIP leverages single-cell RNA sequencing (scRNA-seq) data for its availability, quantitative nature, and genome-wide coverage. It also integrates cell spatial distribution information that we collected using the PhenoCycler^34^ technology, which employs DNA-barcoded antibodies and fluorescence microscopy imaging. While PhenoCycler imaging captures only snapshots of the dynamic bone marrow environment, we used these snapshots to estimate the frequency of cell-cell spatial interactions, assuming that individual cells of the same cell type have similar frequencies of interacting with other cell types.

**Figure 1.**
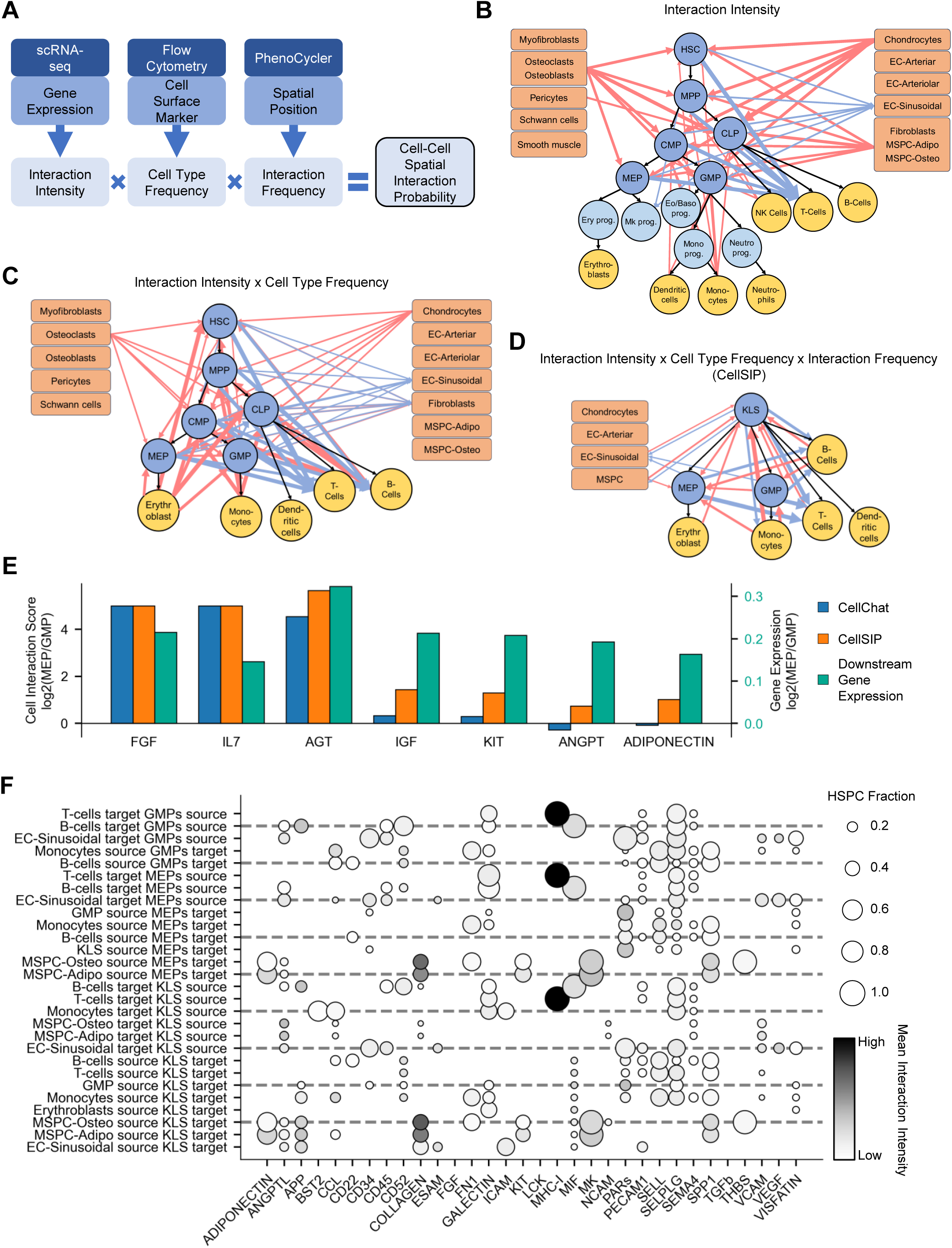
CellSIP quantifies hematopoietic stem and progenitor cell (HSPC) signaling interactions. (A) Data collection and analyses to determine Cell-cell Spatial Interaction Probability (CellSIP). Interaction intensity was calculated based on single-cell RNA sequencing (scRNA-seq) data^33,35^ using CellChat^36^. Cell type frequency in mouse bone marrow was examined by flow cytometry and scRNA-seq analyses (**Tables S1, S2**). Spatial interaction frequency between different cell types was calculated by analyzing PhenoCycler^34^ images (**Tables S3, S4**). (B-D) Cell-cell interaction in the mouse bone marrow between HSPCs (blue), non-hematopoietic cells (orange), and blood and immune cells (yellow). Shown are all bone marrow cell types with available data. Black arrows illustrate the differentiation trajectories of HSPCs. Colored arrows illustrate intercellular signaling interactions. Red arrows illustrate signals targeting HSPCs, and blue arrows illustrate signals sourced by HSPCs. Line thickness reflects interaction strength. (B) Interaction intensity between cell types based on scRNA-seq data. (C) Interaction intensity adjusted by cell type frequency. (D) Cell-cell interaction network based on molecular interaction intensity, cell type frequency, and spatial interaction frequency. (E) Comparing CellChat and CellSIP for intercellular signaling pathways that differentially target MEPs and GMPs. Shown are pathways with significantly different downstream gene expression, which is reflected by either CellChat or CellSIP. (F) Top five signaling pathways mediating each cell-cell interaction illustrated in (D). Color reflects the average interaction intensity. Size indicates the proportion of HSPCs involved in the interaction. Abbreviations: Hematopoietic Stem Cell (HSC), Multipotent Progenitors (MPP), Common Myeloid Progenitors (CMP), Common Lymphoid Progenitors (CLP), Granulocyte-Macrophage Progenitors (GMP), Megakaryocyte-Erythroid Progenitors (MEP), Arterial Endothelial Cells (EC-Arteriar), Arteriolar Endothelial Cells (EC-Arteriolar), Sinusoidal Endothelial Cells (EC-Sinusoidal), Adipogenesis-primed Mesenchymal Stem/Stromal and Progenitor Cells (MSPC-Adipo), and Osteogenesis-primed Mesenchymal Stem/Stromal and Progenitor Cells (MSPC-Osteo). Given the rarity of HSCs, cell-cell spatial interaction was analyzed using cKit^+^Lineage^-^Sca^+^ (KLS) cells that include both HSCs and MPPs. See also Figures S1-S4.

Given that HSPCs have been more rigorously defined and extensively studied in mice than in humans and other organisms, we applied CellSIP to investigate the cell-cell interaction network of HSPCs in mouse bone marrow. We collected scRNA-seq data from six types of HSPCs (n=93,155) in our laboratory, and gathered scRNA-seq data from eighteen types of blood cells^33^ (n=7,497) and thirteen types of non-hematopoietic cells^35^ (n=32,673) from published studies. Leveraging these data sets, we used CellChat^36^ to calculate the molecular interaction intensity between HSPCs and various bone marrow cell types based on their expression levels of ligands and receptors (**Figures 1B and S1A**). The molecular interaction intensity was then adjusted by cell type frequency in mouse bone marrow, as determined by flow cytometry and scRNA-seq analyses (**Figures 1C and S1B; Tables S1, S2**). Finally, we quantified the frequencies of spatial interactions between various cell types using PhenoCycler imaging analyses (**Figure S2; Tables S3, S4**). Given that the effective range of cytokines extends up to 150 µm^37^, we analyzed the overrepresentation and underrepresentation of each bone marrow cell type at various distances within this range from HSPCs. Our analysis revealed that most cell types exhibit stable or peak representation within a 20 µm radius of HSPCs and control random bone marrow cells (**Figure S3**). Therefore, we define the spatial interaction frequency as the average ratio of a cell type’s frequency within 20 µm of a specific type of HSPCs to its overall frequency across the entire tissue section. The spatial interaction frequency was used to further adjust cell-cell interaction intensities, generating the final CellSIP scores (**Figures 1D and S1C; Table S3**).

Our CellSIP analysis begins with 38 bone marrow cell types and over 200 signaling pathways. The spatial interaction analysis is limited to 12 cell types due to constraints in antibody availability and quality. Additionally, HSCs and multipotent progenitors (MPPs) are combined into a cKit+Lineage-Sca+ (KLS) cell population, given the rarity of HSCs and the similar expression of signaling genes between MPPs and HSCs (**Figure S4A**). To validate CellSIP, we analyzed downstream effector gene expression using our scRNA-seq data and the CellCall database (**Figure 1E**).^38^ We identified signaling pathways with significantly different downstream gene expression between megakaryocyte-erythroid progenitors (MEPs) and granulocyte-macrophage progenitors (GMPs). We then compared the signaling interaction intensities of these pathways predicted by CellChat and CellSIP. Our analysis showed that CellSIP predictions more closely align with the corresponding downstream gene expression (**Figure 1E**). These results demonstrate that CellSIP provides more accurate quantifications of cell-cell signaling interactions in dynamic environments such as the bone marrow.

### CellSIP reveals the cell-cell signaling interaction network underlying hematopoiesis

Using CellSIP, we identified interactions between KLS cells and their known niche cells, including adipogenesis-primed mesenchymal stem and progenitor (MSPC-Adipo) cells and sinusoidal endothelial cells (**Figure 1D**)^11,12,39^, as well as known signaling pathways, such as the Kit, MK, Collagen, and Adiponectin pathways, that mediate these cell-cell interactions (**Figure 1F**).^5–9^ Leveraging the quantitative capabilities of CellSIP, we further uncovered the fraction of HSPCs involved, the relative contributions of each bone marrow cell type, and the corresponding interaction intensity. For example, both the Kit and MK pathways are predominantly activated by MSPC-Adipo rather than sinusoidal endothelial cells (**Figure 1F**). The MK pathway exhibits greater intensity and engages a larger portion of KLS cells than the Kit pathway (**Figure 1F**). In contrast, the intercellular adhesion molecule (ICAM) pathway is predominantly mediated by sinusoidal endothelial cells, specifically targeting KLS cells but not oligopotent hematopoietic progenitor cells (**Figure 1F**). These data demonstrate CellSIP’s capacity to identify the primary drivers of the signaling interactions.

CellSIP also uncovered previously overlooked interactions between HSPCs and immune cells, as well as among various types of HSPCs (**Figures 1D, 1F**). The GALECTIN pathway mediates interactions between various types of HSPCs and immune cells. The interactions between HSPCs and T-cells are particularly strong and primarily mediated by the MHC-I pathway (**Figure 1F**). In addition, the protease-activated receptors (PARs) pathway was found to mediate interactions among various types of HSPCs (**Figure 1F**). HSPCs act as sources of signals, including vascular cell adhesion molecule (VCAM) and vascular endothelial growth factor (VEGF) signals, to sinusoidal endothelial cells (**Figure 1F**). VCAM and VEGF have been shown to play roles in HSPC niche homing and survival.^30–32,40^ Altogether, these signaling interactions indicate the presence of feedback mechanisms, contributing to the complex signaling interaction network underlying hematopoiesis.

### Signaling pathway correlations across individual HSPCs

CellSIP analyzes cell-cell signaling interactions among various bone marrow cell types (**Figure 1**), as cell type frequency and spatial interaction frequency were examined at the cell population level. However, within each HSPC population, we observed substantial heterogeneity in intercellular signaling among individual cells, driven by gene expression variability that defines their signaling capacities (**Figure 2A**). Since each individual HSPC is simultaneously engaged in multiple signaling activities, we hypothesized that their engagement in specific signaling pathways could influence their activities in other pathways. Understanding the correlations between signaling pathways could elucidate the signaling cascade in the bone marrow cell network. To test this hypothesis, we calculated the Spearman’s rank correlation between signaling pathways across individual HSCs and identified 411 significant positive correlations and 181 significant negative correlations among 70 signaling pathways in HSCs (**Figures 2A-2B**). For example, HSCs that send more PARs signals are likely to send more CD52 and CCL signals, and fewer ANGPT, MHC-I, and MHC-II signals, as well as to receive fewer APP signals (**Figures 2A-2B**). These signaling pathway correlations involve a variety of interacting cell types (**Figure 2A**), forming a coordinated signaling transduction network in the bone marrow.

**Figure 2.**
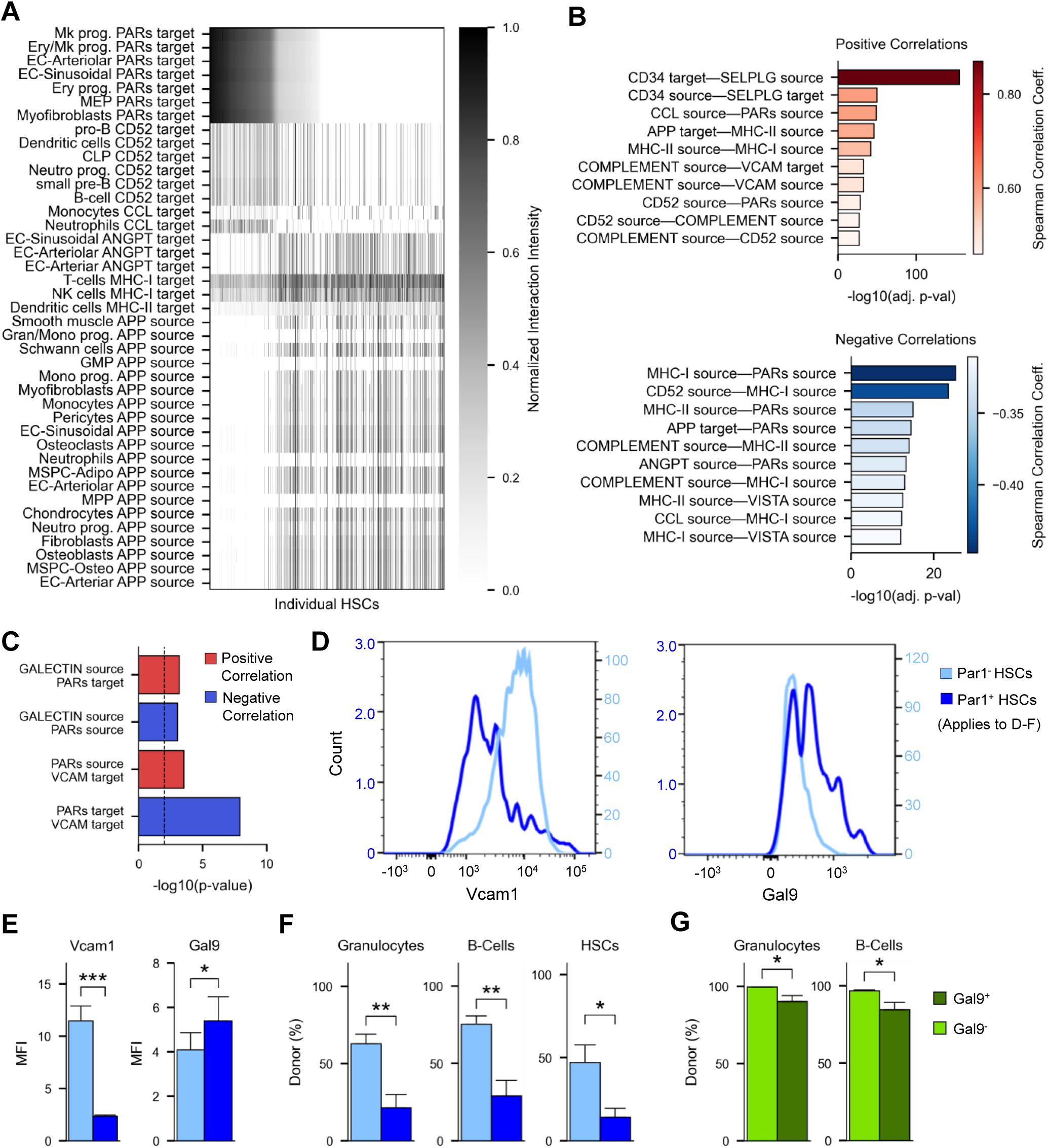
Signaling pathway correlations across individual HSCs. (A) Example signaling pathways exhibiting significant correlations among individual HSCs. Each column represents an individual HSC. HSCs were ordered by their PARs signal intensity sent to megakaryocyte progenitors. (B) Top ten most significant positive and negative pathway correlations across individual HSCs. Common pathways across different cell types were aggregated by averaging Spearman’s rank correlation coefficient and calculating the adjusted p-value using Fisher’s *z-*transformation. (C) PARs pathway significantly correlates with GALECTIN and VCAM pathways. (D) Vcam1 (left) and Gal9 (right) expression on Par1^-^ and Par1^+^ HSCs. The y-axis represents cell count, and the x-axis indicates fluorescence intensity. (E) Median fluorescence intensity (MFI) of Vcam1 (left) and Gal9 (right) expression on Par1^-^ and Par1^+^ HSCs. (B-D) Data derived from flow cytometry analysis of three independent experiments with a total of 6 mice. (F) Equal numbers of Par1^+^ and Par1^-^ HSCs were transplanted into mice (N=5 for the Par1^+^ group; N=8 for the Par1^-^ group). The experiment was performed twice independently. Shown are the donor chimerisms of B-cells and granulocytes in the peripheral blood, as well as HSCs in the bone marrow four months post-transplantation. (G) Equal numbers of Gal9^+^ and Gal9^-^ HSCs were transplanted into mice (N=4 for the Gal9^+^ group; N=5 for the Gal9^-^ group). The experiment was performed twice independently. Shown are the donor chimerisms of B-cells and granulocytes in the peripheral blood four months post-transplantation. (E-H) * p-value < 0.05; ** p-value < 0.01; *** p-value < 0.001. See also Figure S5.

We validated the pathway correlations at the protein level using flow cytometry analysis (**Figures 2C-2E, S5**), focusing specifically on the PARs pathway and its correlation with the VCAM and GALECTIN pathways (**Figure 2C**). The PARs pathway is involved in the most significant pathway correlations in HSCs (**Figure 2B**) and mediates interactions between different types of HSPCs (**Figure 1F**), while the VCAM and GALECTIN pathways play key roles in mediating HSC interactions with sinusoidal endothelial cells and immune cells, respectively (**Figure 1F**). Our results show that Par1^-^ HSCs express significantly higher levels of Vcam1 and significantly lower levels of Galectin-9 (Gal9) on their cell surfaces compared to Par1^+^ HSCs (**Figures 2D-2E, S5**), consistent with our prediction (**Figure 2C**).

### Signaling pathway correlation enables functional prediction

Correlations between signaling pathways suggest that these pathways may share common biological functions. This is exemplified by the validated correlations between the PARs, VCAM, and GALECTIN pathways (**Figures 2C-2E, S5**). The VCAM pathway has been shown to regulate HSPC homing to their bone marrow niche.^30–32^ Similarly, the PARs signaling pathway has been linked to the bone marrow retention of HSCs through the EPCR and CXCL12 pathways.^41,42^ However, functional differences between HSCs differentially engaged in the PARs signaling pathway haven’t been investigated. Moreover, the role of GALECTIN signaling in HSC regulation remains unexplored.

To demonstrate the utility of signaling pathway correlations in predicting the functional impact of signaling pathways, we conducted transplantation experiments using equal numbers of Par1^+^ and Par1^-^ HSCs, as well as Gal9^+^ and Gal9^-^ HSCs, each co-transplanted with identical numbers of unfractionated bone marrow cells (**Figures 2F-2G**). Four months post-transplantation, Par1^-^ HSCs showed significantly higher levels of hematopoietic reconstitution compared to Par1^+^ HSCs (**Figure 2F**). Similarly, Gal9^-^ HSCs produced significantly greater hematopoietic reconstitution than Gal9^+^ HSCs (**Figure 2G**). This suggests that correlated signaling pathways have a similar functional impact.

### Signaling pathway correlations shift during hematopoiesis

Comparing signaling pathway correlations across different types of HSPCs, we found that hematopoietic progenitors involved in myeloid differentiation, such as MPPs, CMPs, and GMPs, exhibit relatively low numbers of pathway correlations, while HSCs and common lymphoid progenitors (CLPs) show the highest numbers of correlations (**Figure 3A**). This suggests that the dynamics of intercellular communication evolve over the course of hematopoiesis and differ between various lineages. In general, positive correlations between signaling pathways are stronger and more consistently maintained throughout hematopoiesis, indicating a stable cooperative mechanism essential throughout hematopoiesis (**Figure 3B**). In contrast, negative correlations tend to emerge at distinct hematopoietic stages, suggesting that antagonistic signaling interactions play crucial roles in regulating specific phases of hematopoiesis (**Figure 3B**).

**Figure 3.**
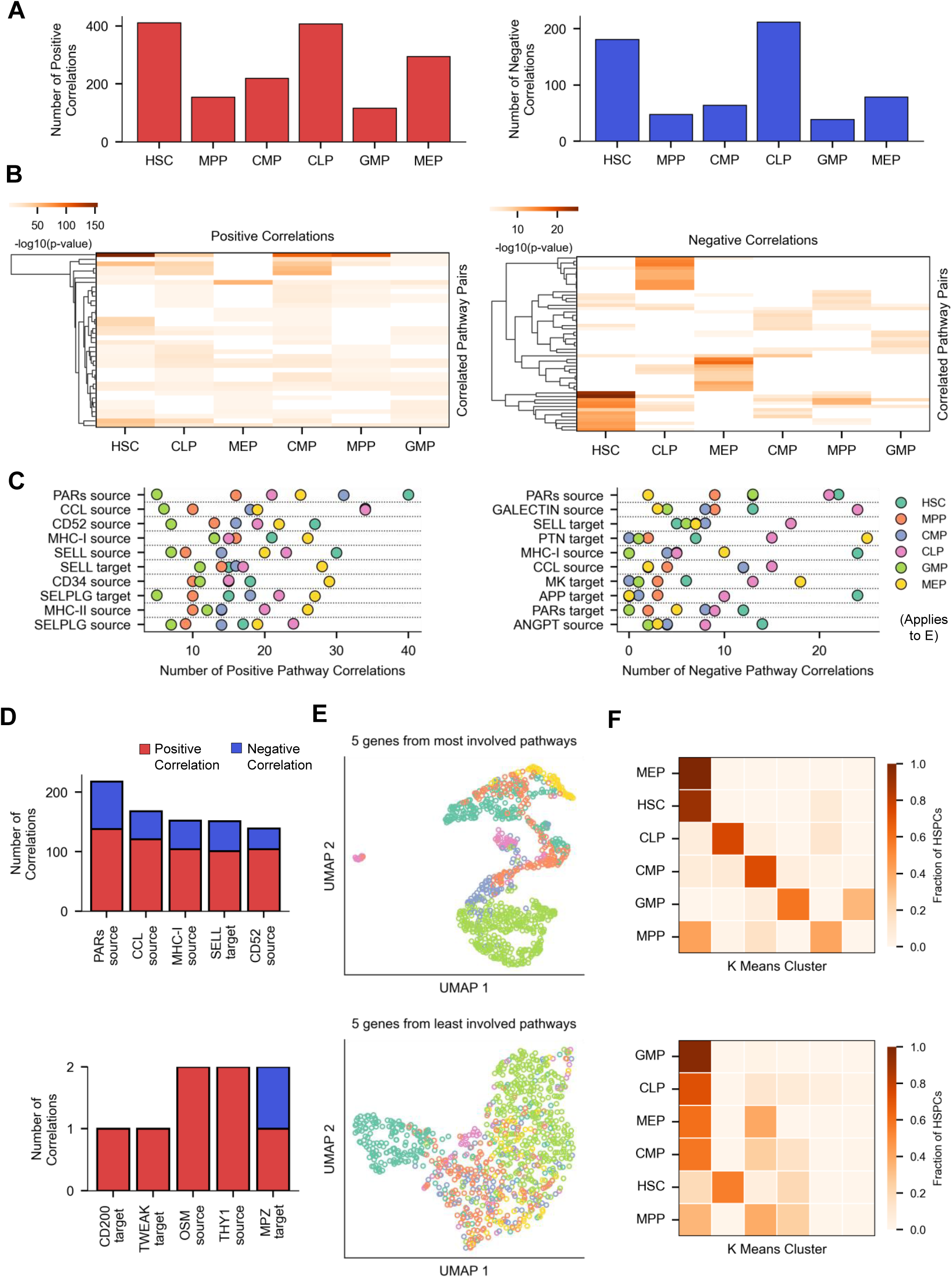
Signaling pathway correlations in different types of HSPCs. (A) Number of significantly correlated signaling pathway pairs identified in each type of HSPCs. (B) Comparing signaling pathway correlations across different types of HSPCs. Rows show the top ten most significant positive and negative correlations in each type of HSPCs. Color reflects the p-value of the Spearman’s rank correlation. White represents adjusted p-value > 0.001. (C) Signaling pathways correlate with different numbers of other pathways in different types of HSPCs. Shown are the top ten signaling pathways most frequently involved in positive and negative correlations across various types of HSPCs. (D) Signaling pathways most (top) or least (bottom) frequently involved in pathway correlations among HSPCs. (E) UMAP visualization of HSPC single-cell RNA-seq data, showing metacells clustered by 5 highest expressed genes from 5 signaling pathways that are most (top) or least (bottom) frequently involved in pathway correlations as shown in (D) (**Tables S5, S6**). (F) K means clustering (k=6) using 5 highest expressed genes from 5 signaling pathways that are most (top) or least (bottom) frequently involved in pathway correlations as shown in (D) (**Tables S5, S6**). Each column represents a cluster. Each row represents one type of HSPCs (abbreviations provided in the Figure 1 legend). Heatmap colors indicate the fraction of HSPCs in each cluster. See also Figures S6,S7.

Among the top ten signaling pathways that are most frequently involved in pathway correlations, each pathway correlates with varying numbers of other pathways in different types of HSPCs (**Figure 3C**). Moreover, a single signaling pathway can correlate with different pathways in different types of HSPCs (**Figure S6**). For example, the PARs signaling pathway positively correlates with the VCAM pathways exclusively in HSCs, with the TGF-β and PECAM1 pathways exclusively in MEPs, and negatively correlates with the IL7 pathway exclusively in CLPs (**Figure S6**). These data further illustrate the specificity of signaling pathway correlations at distinct stages of hematopoiesis. To determine how these pathway correlations contribute to distinct hematopoietic stages, we identified the signaling pathways most frequently involved in positive and negative correlations (**Figure 3D**). Remarkably, as few as five genes from the top five signaling pathways can effectively cluster individual HSPCs into groups that match their conventional cell types (**Figures 3E**, **3F**, **S7**, **Tables S5**, **S6**). This highlights the utility of signaling pathway correlations in identifying biomarkers and regulators for distinct hematopoietic stages.

### Hematopoiesis Intercellular Signaling Explorer (HISE)

Our study mapped and quantified the signaling interactions between mouse HSPCs and various bone marrow cell types, as well as their signaling pathway correlations (**Figures 1-3**). To provide access to this comprehensive dataset, we developed the Hematopoiesis Intercellular Signaling Explorer, a user-friendly and interactive database offering a systems-level understanding of cell-cell communication in mouse bone marrow hematopoiesis.

HISE is organized by bone marrow cell types and signaling pathways (**Figure 4A**). Users can explore specific cell types or pathways of interest to obtain information including the fractions of HSPCs engaged in the pathways, bone marrow cell types involved, intensity of pathway interactions, pathway correlations, and genes associated with each pathway (**Figure 4A**). For example, when users search for the interactions between a specific type of HSPCs and a bone marrow cell type, such as HSCs and myofibroblasts, HISE displays the potential signaling pathways that mediate their interactions (**Figure 4B**). Similarly, when users search for a specific signaling pathway, such as the APP pathway in HSCs, HISE displays the bone marrow cell types involved (**Figure 4C**). Moreover, HISE provides detailed information on the fraction of HSCs involved in these interactions and the intensity of their interactions with each bone marrow cell type (**Figures 4B-4C**). On the pathway correlation page of HISE, pathways are presented in a network graph that illustrates their positive and negative correlations in a select type of HSPC (**Figure 4D**). Users can customize the display by setting independent thresholds for the correlation coefficient of positive and negative correlations. Lastly, HISE provides a gene network diagram for each signaling pathway that features ligands and receptors involved, as well as their potential interactions (**Figure 4E**). All data are available for download to facilitate further analysis.

**Figure 4.**
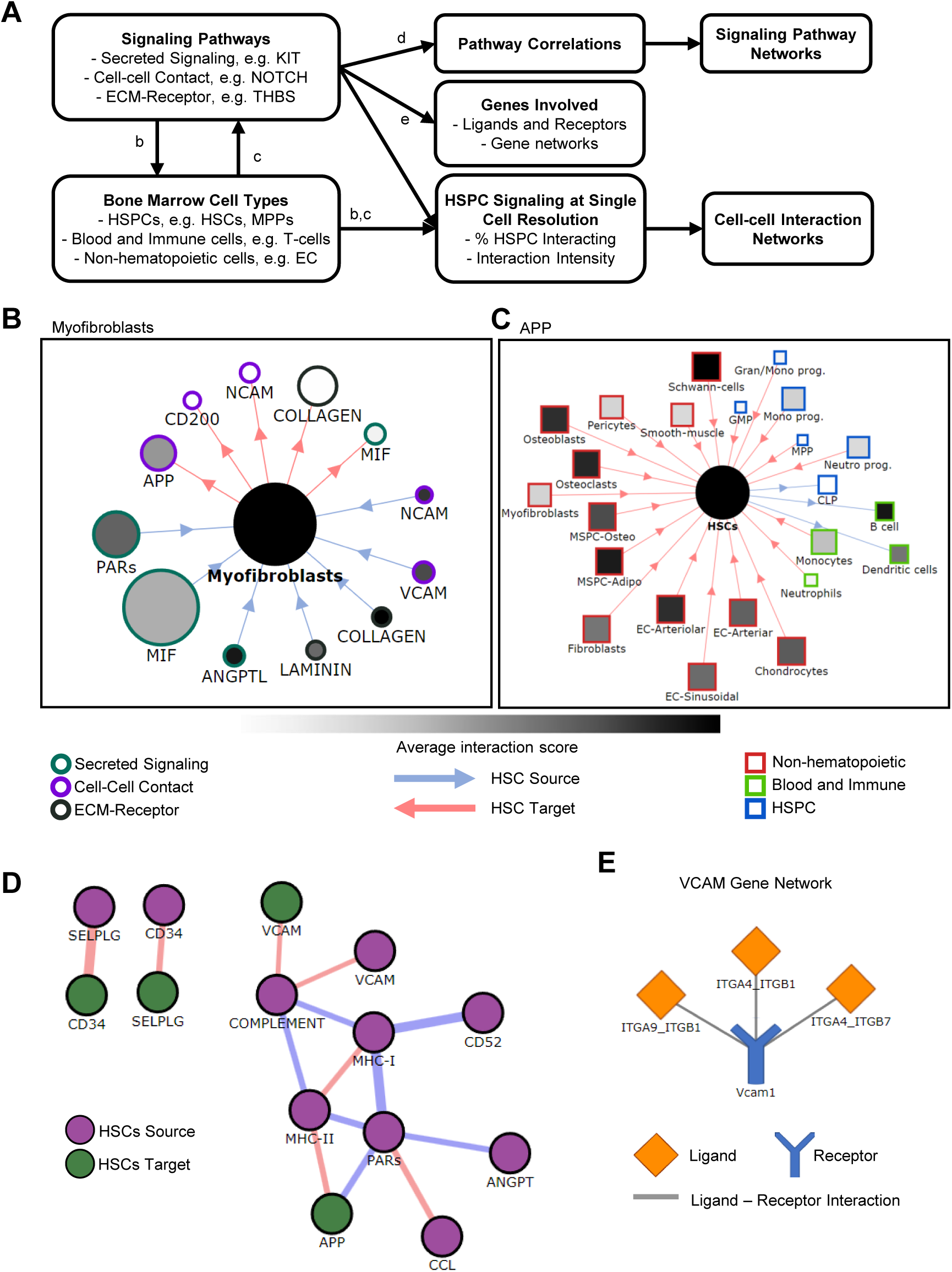
Hematopoiesis Intercellular Signaling Explorer (HISE). (A) Database components of HISE. For each type of HSPC, HISE offers detailed information on relevant signaling pathways and associated bone marrow cell types. HISE also provides data about signaling pathway correlations, involved genes, proportion of HSPCs engaged, and interaction intensity. (B-E) Sample visualizations from HISE. (B) Sample output from HISE: searching for HSCs and myofibroblasts as an example. Each dot represents a signaling pathway that mediates the interactions between HSCs and myofibroblasts. The color of each dot depicts the interaction intensity, while the size indicates the proportion of HSCs involved in each pathway. (C) Sample output from HISE: searching for HSCs and the APP pathway as an example. Each square marker represents a bone marrow cell type that can interact with HSCs through the APP pathway. The color of each marker depicts the interaction intensity, while the size indicates the proportion of the bone marrow cells involved. (D) Sample pathway correlation network of HSCs from HISE. Each dot represents a signaling pathway involving HSCs. Edges between dots indicate pathway correlations (red: positive; blue: negative), with edge width indicating correlation strength. (E) Sample gene network plot from HISE that displays ligands, receptors, and their interactions.

## DISCUSSION

In this study, we investigated the cellular and molecular networks of intercellular signaling in bone marrow hematopoiesis by quantifying the signaling intensity, identifying the key cellular and molecular components, and uncovering the signaling correlations in HSPCs. We generated a comprehensive database for exploring cell-cell interactions and signaling transductions in the mouse bone marrow, which is accessible through HISE (**Figure 4**). Researchers can utilize HISE to learn about potential signaling transductions relevant to their specific cell types of interest, fostering the development of new hypotheses about cell fate regulation. They can also identify key genes and cell types involved in specific signaling pathways, which can help pinpoint critical contributors and examine their regulatory mechanisms. Given the complexity of bone marrow cell interactions and the resilience of blood and immune cell regeneration, a systems-level understanding of HSPC communication is crucial for understanding hematopoiesis. Similar approaches can be applied to studying human hematopoiesis, although human HSPCs remain poorly defined and characterized due to genetic variability across individuals. Here, using a well-characterized mouse strain and comprehensive scRNA-seq data from several laboratories including our own, we demonstrate the potential of HISE in providing insights into the regulation of the hematopoietic system.

To quantify cell-cell signaling interactions, we developed CellSIP that takes into account the semi-fluidic nature of the bone marrow (**Figure 1**). CellSIP provides quantification of the signaling intensity and the relative contribution of each cell type, enabling researchers to identify the dominant regulators. This algorithm incorporates cell-cell spatial interaction data, illustrating how spatial measurements can be integrated with intercellular signaling analysis. In addition, CellSIP accounts for the frequency of each cell type in the bone marrow to calculate its contribution to signaling transduction. It is important to note that while signaling interactions primarily take place at the protein level, CellSIP is based on scRNA-seq data, which has limitations such as low sensitivity and a high dropout rate. Moreover, protein and mRNA levels may differ for some genes. Nonetheless, scRNA-seq data allows for a genome-wide profile of potential signaling interactions across all major cell types. Moreover, mRNA-level analyses can reveal insights into biological processes that are not evident at the protein level. CellSIP is designed for studying cell-cell signaling in the bone marrow, where cell migration is constant. It can also be applied to other biological processes characterized by prevalent cell migration, such as in development and tissue regeneration.

Our study reveals signaling pathway correlations among individual HSPCs at the same stage of hematopoietic differentiation by leveraging scRNA-seq data and CellChat (**Figures 2-3**). These correlations indicate that one pathway may regulate another, or that both pathways may share a common upstream regulator. The identified pathway correlations highlight the inherent heterogeneity of HSPC signaling and provide insights into the organization of cellular and signaling networks underlying hematopoiesis. We show that the pathway correlations can distinguish different stages of hematopoiesis (**Figures 3E, 3F, S7**), and can also help identify new regulators through their associations with established ones, as illustrated by the PARs, GALECTIN, and VCAM pathway correlations (**Figures 2F-2G**). Notably, the PARs signaling pathway is the most frequently involved in pathway correlations among HSPCs (**Figure 3D**), found in both negative and positive correlations across individual HSCs (**Figure 3C**), as well as in communications between different types of HSPCs (**Figure 1F**). Our transplantation experiments with Par1^+^ and Par1^-^ HSCs demonstrated the crucial role of the PARs signaling in regulating hematopoiesis (**Figure 2F**). Altogether, these findings reveal the pivotal role of the PARs pathway in coordinating hematopoiesis. In the future, the signaling pathway correlation analysis can be extended to other tissues and organs to explore intercellular signaling networks and identify key regulators.

**Supplemental Figure 1.**
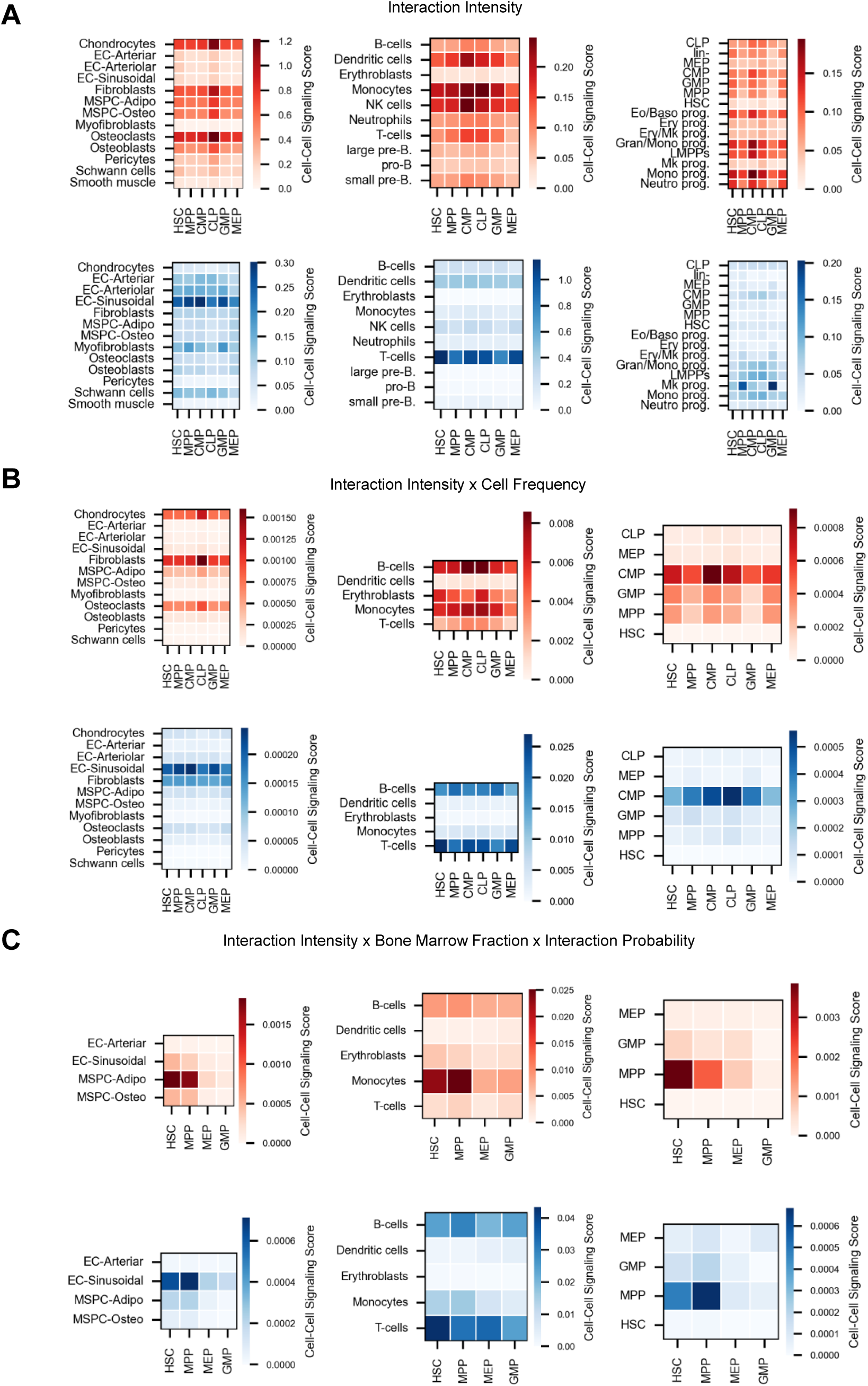
Quantification of cell-cell signaling interactions in mouse bone marrow, related to Figure 1. (A) Interaction intensity between cell types based on scRNA-seq data and CellChat^36^. (B) Interaction intensity adjusted by cell type frequency. (C) Final cell-cell signaling scores integrating data from molecular interaction intensity, cell type frequency, and spatial interaction frequency. Red illustrates signals targeting HSPCs. Blue illustrates signals sourced by HSPCs. Some bone marrow cell types are excluded due to lack of data.

**Supplemental Figure 2.**
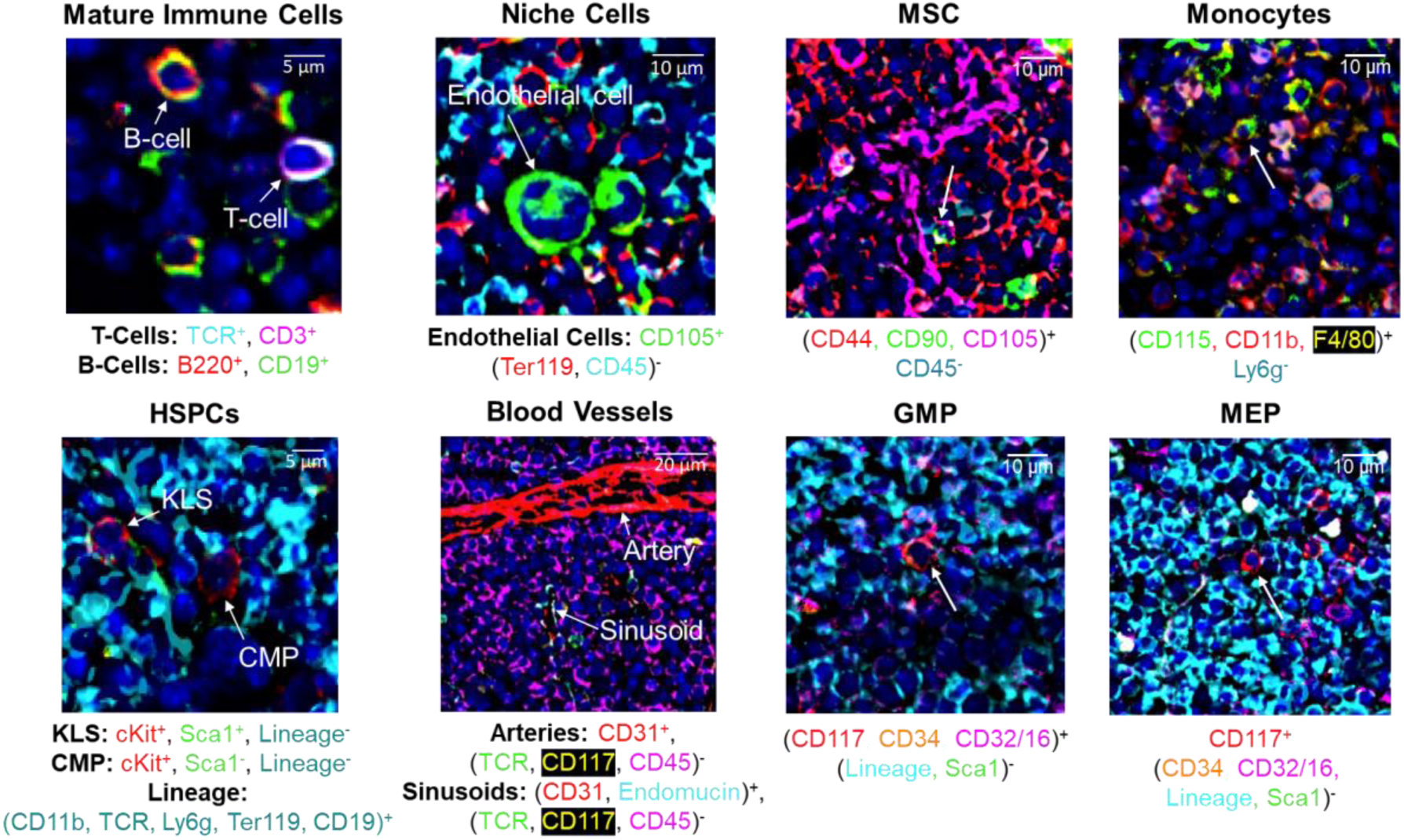
PhenoCycler images used to determine cell-cell spatial interactions, related to Figure 1. Representative images from PhenoyCycler analysis showing antibody stains and cell type annotations. Please refer to the Figure 1 legend for abbreviations.

**Supplemental Figure 3.**
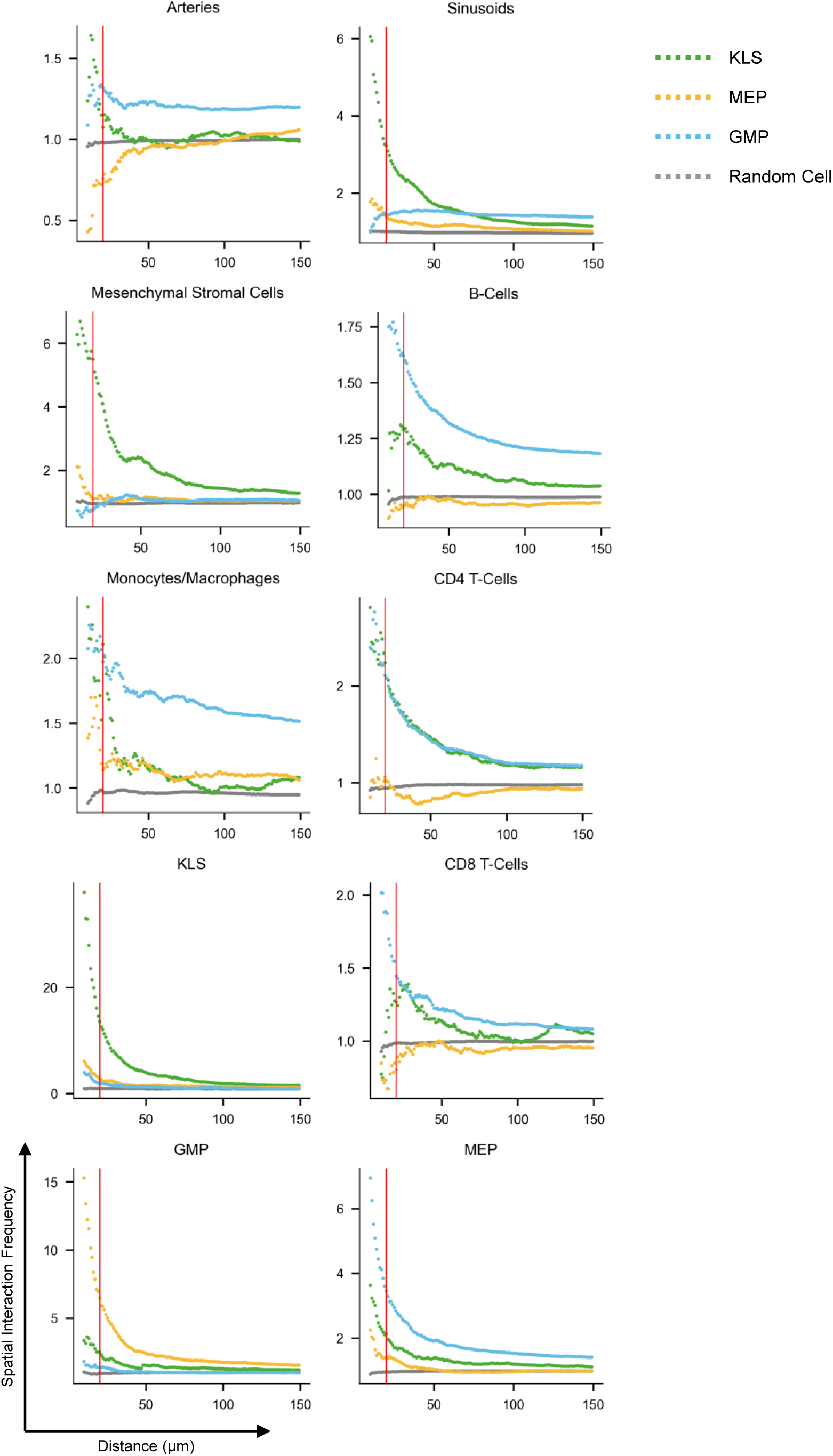
Quantifying spatial interactions between HSPCs and bone marrow cells, related to Figure 1. Quantification of spatial interaction frequency between bone marrow cells and HSPCs, or random bone marrow cells as control. The spatial interaction frequency (y-axis) is calculated as the average ratio of a cell type’s frequency within a specified radius (x-axis) of HSPCs to its overall frequency throughout the entire tissue section. Please refer to the Figure 1 legend for abbreviations.

**Supplemental Figure 4.**
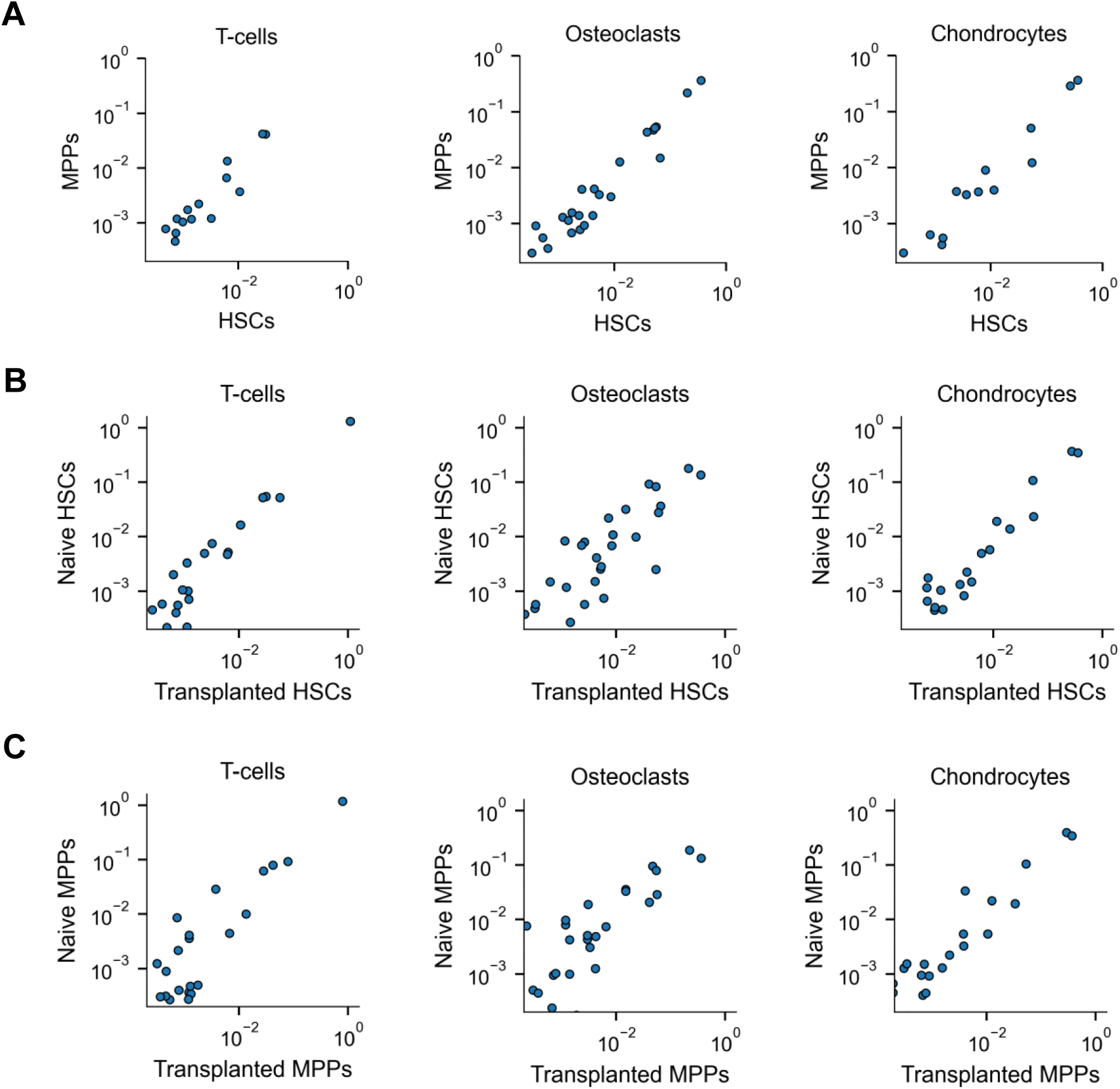
Comparison of HSPC intercellular signaling intensity, related to Figure 1. Each dot represents a signaling pathway. Shown is the average interaction intensity of individual HSCs and MPPs with three bone marrow cell types that exhibit the strongest interactions with HSCs. (A) Comparison of signaling interaction intensity between HSCs and MPPs. (B) Comparison of HSC signaling interaction intensity between naive and transplanted mice. (C) Comparison of MPP signaling interaction intensity between naive and transplanted mice.

**Supplemental Figure 5.**
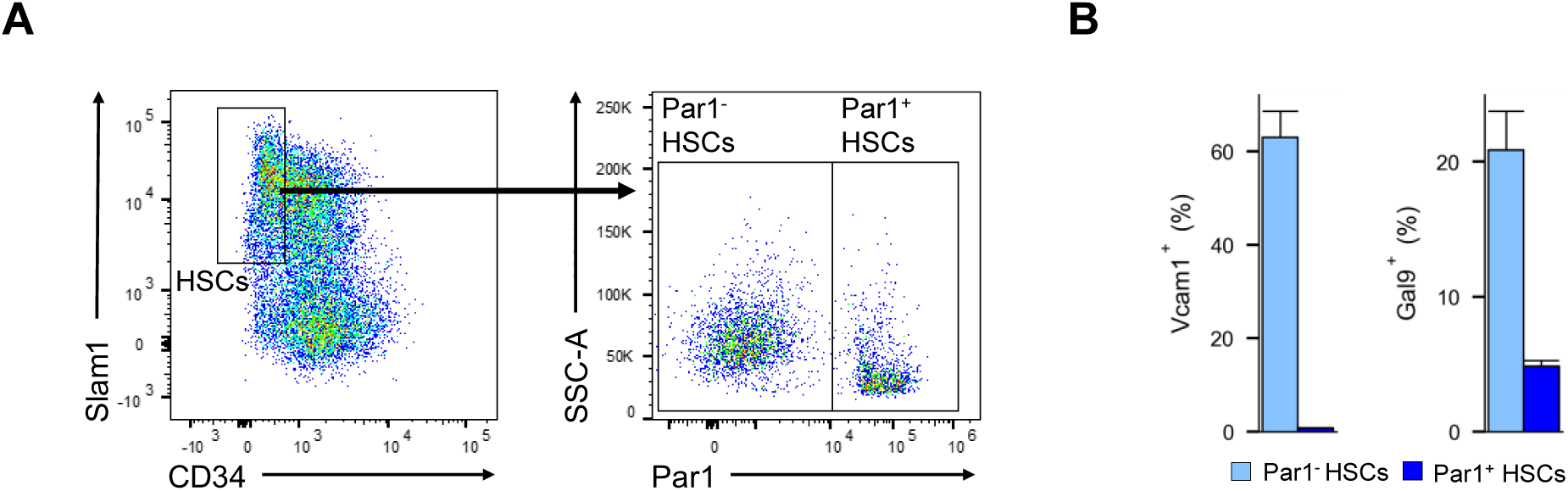
Flow cytometry analysis of Par1^+^ HSCs and Par1^-^ HSCs, related to Figure 2. (A) Gating of Par1^+^ HSCs and Par1^-^ HSCs. Shown are mouse bone marrow cells pre-gated for cKit^+^Lineage^-^ Sca^+^Flk2^-^. (B) Fraction of Vcam1^+^ or Gal9^+^ cells among Par1^+^ HSCs and Par1^-^ HSCs.

**Supplemental Figure 6.**
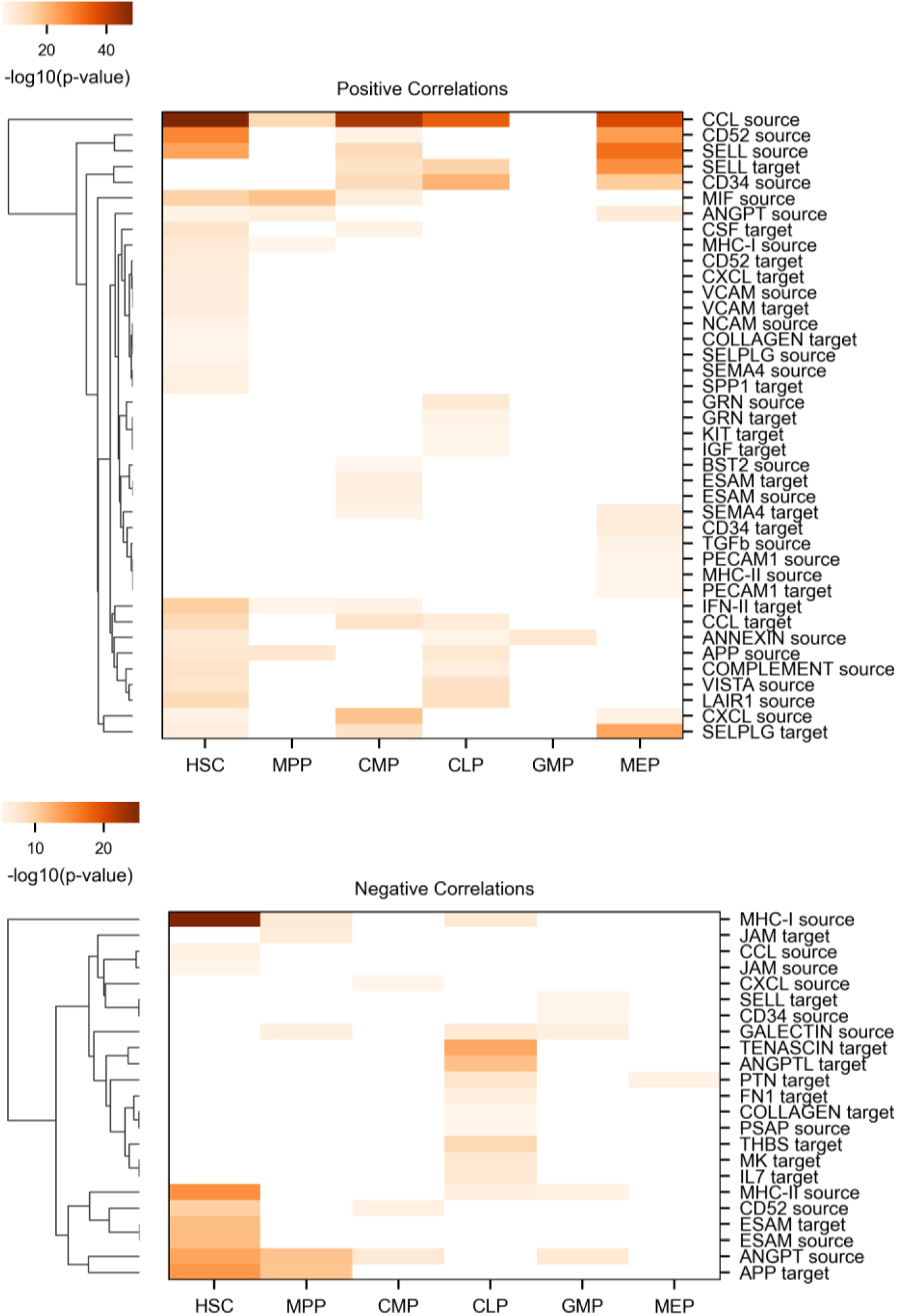
Signaling pathways significantly correlated with the PARs pathway in various types of HSPCs, related to Figure 3. Color reflects the p-value of the Spearman’s rank correlation. White represents adjusted p-value > 0.001.

**Supplemental Figure 7.**
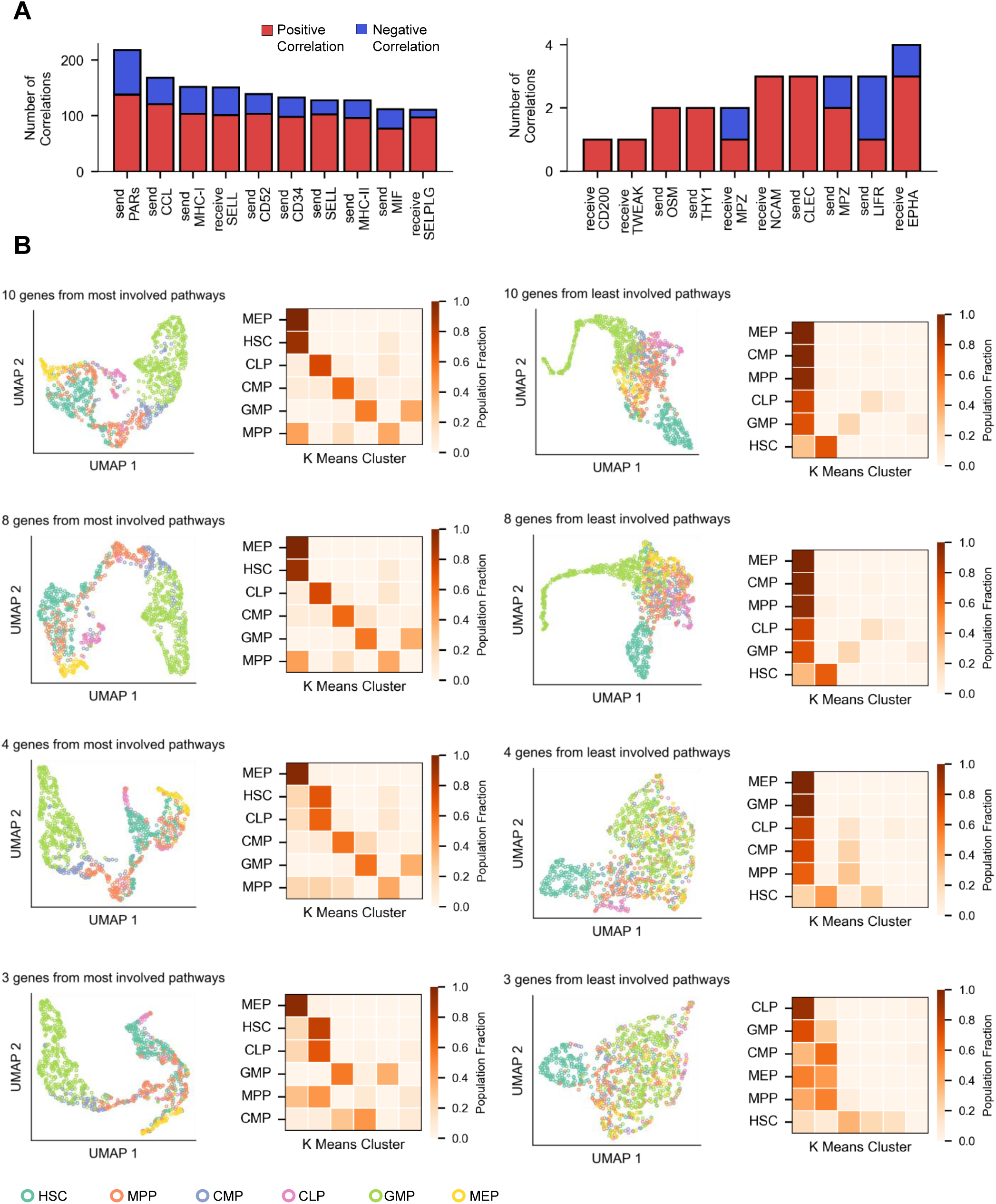
A few signaling pathway genes effectively cluster HSPCs, related to Figure 3. (A) Signaling pathways that are most (left) or least (right) frequently involved in pathway correlations. (B) UMAP visualization of HSPC scRNA-seq data, showing metacells clustered by various numbers of genes from signaling pathways that are most (left) or least (right) frequently involved in pathway correlations as shown in (A) (Gene names provided in **Tables S5, S6** respectively). K means clustering (k=6) using various numbers of genes from signaling pathways that are most (left) or least (right) frequently involved in pathway correlations as shown in (A) (Gene names provided in **Tables S5, S6** respectively). Each column represents a cluster. Each row represents one type of HSPCs (abbreviations provided in the Figure 1 legend). Heatmap colors indicate the fraction of HSPCs in each cluster.

## References

1. Weissman, I.L., and Shizuru, J.A. (2008). The origins of the identification and isolation of hematopoietic stem cells, and their capability to induce donor-specific transplantation tolerance and treat autoimmune diseases. Blood 112, 3543–3553. 10.1182/blood-2008-08-078220.

2. Seita, J., and Weissman, I.L. (2010). Hematopoietic stem cell: self-renewal versus differentiation. WIREs Syst. Biol. Med. 2, 640–653. 10.1002/wsbm.86.

3. Wilson, A., Laurenti, E., Oser, G., Wath, R.C. van der, Blanco-Bose, W., Jaworski, M., Offner, S., Dunant, C.F., Eshkind, L., Bockamp, E., et al. (2008). Hematopoietic Stem Cells Reversibly Switch from Dormancy to Self-Renewal during Homeostasis and Repair. Cell 135, 1118–1129. 10.1016/j.cell.2008.10.048.

4. Pietras, E.M. (2017). Inflammation: a key regulator of hematopoietic stem cell fate in health and disease. Blood 130, 1693–1698. 10.1182/blood-2017-06-780882.

5. Thorén, L.A., Liuba, K., Bryder, D., Nygren, J.M., Jensen, C.T., Qian, H., Antonchuk, J., and Jacobsen, S.-E.W. (2008). Kit Regulates Maintenance of Quiescent Hematopoietic Stem Cells. J. Immunol. 180, 2045–2053. 10.4049/jimmunol.180.4.2045.

6. Sorrelle, N., Dominguez, A.T.A., and Brekken, R.A. (2017). From top to bottom: midkine and pleiotrophin as emerging players in immune regulation. J. Leukoc. Biol. 102, 277–286. 10.1189/jlb.3MR1116-475R.

7. Klein, G. (1995). The extracellular matrix of the hematopoietic microenvironment. Experientia 51, 914–926. 10.1007/BF01921741.

8. Çelebi, B., Pineault, N., and Mantovani, D. (2012). The Role of Collagen Type I on Hematopoietic and Mesenchymal Stem Cells Expansion and Differentiation. Adv. Mater. Res. 409, 111–116. 10.4028/www.scientific.net/AMR.409.111.

9. Meacham, C.E., Jeffery, E.C., Burgess, R.J., Sivakumar, C.D., Arora, M.A., Stanley, A.M., Colby, E.M., Crane, G.M., Zhao, Z., and Morrison, S.J. (2022). Adiponectin receptors sustain haematopoietic stem cells throughout adulthood by protecting them from inflammation. Nat. Cell Biol. 24, 697–707. 10.1038/s41556-022-00909-9.

10. Chan, C.K.F., Chen, C.-C., Luppen, C.A., Kim, J.-B., DeBoer, A.T., Wei, K., Helms, J.A., Kuo, C.J., Kraft, D.L., and Weissman, I.L. (2009). Endochondral ossification is required for haematopoietic stem-cell niche formation. Nature 457, 490–494. 10.1038/nature07547.

11. Aqmasheh, S., Shamsasanjan, karim, Akbarzadehlaleh, P., Pashoutan Sarvar, D., and Timari, H. (2017). Effects of Mesenchymal Stem Cell Derivatives on Hematopoiesis and Hematopoietic Stem Cells. Adv. Pharm. Bull. 7, 165–177. 10.15171/apb.2017.021.

12. Greenbaum, A., Hsu, Y.-M.S., Day, R.B., Schuettpelz, L.G., Christopher, M.J., Borgerding, J.N., Nagasawa, T., and Link, D.C. (2013). CXCL12 in early mesenchymal progenitors is required for haematopoietic stem-cell maintenance. Nature 495, 227–230. 10.1038/nature11926.

13. Butler, J.M., Nolan, D.J., Vertes, E.L., Varnum-Finney, B., Kobayashi, H., Hooper, A.T., Seandel, M., Shido, K., White, I.A., Kobayashi, M., et al. (2010). Endothelial cells are essential for the self-renewal and repopulation of Notch-dependent hematopoietic stem cells. Cell Stem Cell 6, 251–264. 10.1016/j.stem.2010.02.001.

14. Ding, L., Saunders, T.L., Enikolopov, G., and Morrison, S.J. (2012). Endothelial and perivascular cells maintain haematopoietic stem cells. Nature 481, 457–462. 10.1038/nature10783.

15. Gillette, J.M., Larochelle, A., Dunbar, C.E., and Lippincott-Schwartz, J. (2009). Intercellular transfer to signalling endosomes regulates an ex vivo bone marrow niche. Nat. Cell Biol. 11, 303–311. 10.1038/ncb1838.

16. Zhou, B.O., Ding, L., and Morrison, S.J. (2015). Hematopoietic stem and progenitor cells regulate the regeneration of their niche by secreting Angiopoietin-1. eLife 4, e05521. 10.7554/eLife.05521.

17. Broxmeyer, H.E. (2008). Chemokines in hematopoiesis. Curr. Opin. Hematol. 15, 49–58. 10.1097/MOH.0b013e3282f29012.

18. Blank, U., Karlsson, G., and Karlsson, S. (2008). Signaling pathways governing stem-cell fate. Blood 111, 492–503. 10.1182/blood-2007-07-075168.

19. Klumpe, H.E., Langley, M.A., Linton, J.M., Su, C.J., Antebi, Y.E., and Elowitz, M.B. (2022). The context-dependent, combinatorial logic of BMP signaling. Cell Syst. 13, 388–407.e10. 10.1016/j.cels.2022.03.002.

20. Su, C.J., Murugan, A., Linton, J.M., Yeluri, A., Bois, J., Klumpe, H., Langley, M.A., Antebi, Y.E., and Elowitz, M.B. (2022). Ligand-receptor promiscuity enables cellular addressing. Cell Syst. 13, 408–425.e12. 10.1016/j.cels.2022.03.001.

21. Granados, A.A., Kanrar, N., and Elowitz, M.B. (2024). Combinatorial expression motifs in signaling pathways. Cell Genomics 4, 100463. 10.1016/j.xgen.2023.100463.

22. Antebi, Y.E., Linton, J.M., Klumpe, H., Bintu, B., Gong, M., Su, C., McCardell, R., and Elowitz, M.B. (2017). Combinatorial Signal Perception in the BMP Pathway. Cell 170, 1184–1196.e24. 10.1016/j.cell.2017.08.015.

23. Nombela-Arrieta, C., and Manz, M.G. (2017). Quantification and three-dimensional microanatomical organization of the bone marrow. Blood Adv. 1, 407–416. 10.1182/bloodadvances.2016003194.

24. Upadhaya, S., Krichevsky, O., Akhmetzyanova, I., Sawai, C.M., Fooksman, D.R., and Reizis, B. (2020). Intravital Imaging Reveals Motility of Adult Hematopoietic Stem Cells in the Bone Marrow Niche. Cell Stem Cell 27, 336–345.e4. 10.1016/j.stem.2020.06.003.

25. Lo Celso, C., Fleming, H.E., Wu, J.W., Zhao, C.X., Miake-Lye, S., Fujisaki, J., Côté, D., Rowe, D.W., Lin, C.P., and Scadden, D.T. (2009). Live-animal tracking of individual haematopoietic stem/progenitor cells in their niche. Nature 457, 92. 10.1038/nature07434.

26. Gurkan, U.A., and Akkus, O. (2008). The Mechanical Environment of Bone Marrow: A Review. Ann. Biomed. Eng. 36, 1978–1991. 10.1007/s10439-008-9577-x.

27. Hoggatt, J., and Pelus, L.M. (2011). Mobilization of hematopoietic stem cells from the bone marrow niche to the blood compartment. Stem Cell Res. Ther. 2, 13. 10.1186/scrt54.

28. Winkler, I.G., Sims, N.A., Pettit, A.R., Barbier, V., Nowlan, B., Helwani, F., Poulton, I.J., van Rooijen, N., Alexander, K.A., Raggatt, L.J., et al. (2010). Bone marrow macrophages maintain hematopoietic stem cell (HSC) niches and their depletion mobilizes HSCs. Blood 116, 4815–4828. 10.1182/blood-2009-11-253534.

29. Christodoulou, C., Spencer, J., Yeh, S., Turcotte, R., Kokkaliaris, K., Panero, R., Ramos, A., Guo, G., Seyedhassantehrani, N., Esipova, T., et al. (2020). Live-animal imaging of native hematopoietic stem and progenitor cells. Nature 578, 278–283. 10.1038/s41586-020-1971-z.

30. Stanger, A.M.P., and Lengerke, C. (2022). VCAM1 as a don’t-eat-me molecule. Nat. Cell Biol. 24, 282–283. 10.1038/s41556-022-00864-5.

31. Pinho, S., Wei, Q., Maryanovich, M., Zhang, D., Balandrán, J.C., Pierce, H., Nakahara, F., Di Staulo, A., Bartholdy, B.A., Xu, J., et al. (2022). VCAM1 confers innate immune tolerance on haematopoietic and leukaemic stem cells. Nat. Cell Biol. 24, 290–298. 10.1038/s41556-022-00849-4.

32. Li, D., Xue, W., Li, M., Dong, M., Wang, J., Wang, X., Li, X., Chen, K., Zhang, W., Wu, S., et al. (2018). VCAM-1+ macrophages guide the homing of HSPCs to a vascular niche. Nature 564, 119–124. 10.1038/s41586-018-0709-7.

33. Baccin, C., Al-Sabah, J., Velten, L., Helbling, P.M., Grünschläger, F., Hernández-Malmierca, P., Nombela-Arrieta, C., Steinmetz, L.M., Trumpp, A., and Haas, S. (2020). Combined single-cell and spatial transcriptomics reveal the molecular, cellular and spatial bone marrow niche organization. Nat. Cell Biol. 22, 38–48. 10.1038/s41556-019-0439-6.

34. Goltsev, Y., Samusik, N., Kennedy-Darling, J., Bhate, S., Hale, M., Vazquez, G., Black, S., and Nolan, G.P. (2018). Deep Profiling of Mouse Splenic Architecture with CODEX Multiplexed Imaging. Cell 174, 968–981.e15. 10.1016/j.cell.2018.07.010.

35. Dolgalev, I., and Tikhonova, A.N. (2021). Connecting the Dots: Resolving the Bone Marrow Niche Heterogeneity. Front. Cell Dev. Biol. 9.

36. Jin, S., Guerrero-Juarez, C.F., Zhang, L., Chang, I., Ramos, R., Kuan, C.-H., Myung, P., Plikus, M.V., and Nie, Q. (2021). Inference and analysis of cell-cell communication using CellChat. Nat. Commun. 12, 1088. 10.1038/s41467-021-21246-9.

37. Oyler-Yaniv, A., Oyler-Yaniv, J., Whitlock, B.M., Liu, Z., Germain, R.N., Huse, M., Altan-Bonnet, G., and Krichevsky, O. (2017). A Tunable Diffusion-Consumption Mechanism of Cytokine Propagation Enables Plasticity in Cell-to-Cell Communication in the Immune System. Immunity 46, 609–620. 10.1016/j.immuni.2017.03.011.

38. Zhang, Y., Liu, T., Hu, X., Wang, M., Wang, J., Zou, B., Tan, P., Cui, T., Dou, Y., Ning, L., et al. (2021). CellCall: integrating paired ligand–receptor and transcription factor activities for cell–cell communication. Nucleic Acids Res. 49, 8520–8534. 10.1093/nar/gkab638.

39. Sacchetti, B., Funari, A., Michienzi, S., Di Cesare, S., Piersanti, S., Saggio, I., Tagliafico, E., Ferrari, S., Robey, P.G., Riminucci, M., et al. (2007). Self-renewing osteoprogenitors in bone marrow sinusoids can organize a hematopoietic microenvironment. Cell 131, 324–336. 10.1016/j.cell.2007.08.025.

40. Gerber, H.-P., and Ferrara, N. (2003). The role of VEGF in normal and neoplastic hematopoiesis. J. Mol. Med. Berl. Ger. 81, 20–31. 10.1007/s00109-002-0397-4.

41. Gur-Cohen, S., Itkin, T., Chakrabarty, S., Graf, C., Kollet, O., Ludin, A., Golan, K., Kalinkovich, A., Ledergor, G., Wong, E., et al. (2015). PAR1 signaling regulates the retention and recruitment of EPCR-expressing bone marrow hematopoietic stem cells. Nat. Med. 21, 1307–1317. 10.1038/nm.3960.

42. Gur-Cohen, S., Kollet, O., Graf, C., Esmon, C.T., Ruf, W., and Lapidot, T. (2016). Regulation of long-term repopulating hematopoietic stem cells by EPCR/PAR1 signaling. Ann. N. Y. Acad. Sci. 1370, 65–81. 10.1111/nyas.13013.

43. Brewer, C., Chu, E., Chin, M., and Lu, R. (2016). Transplantation Dose Alters the Differentiation Program of Hematopoietic Stem Cells. Cell Rep. 15, 1848–1857. 10.1016/j.celrep.2016.04.061.

44. Nguyen, L., Wang, Z., Chowdhury, A.Y., Chu, E., Eerdeng, J., Jiang, D., and Lu, R. (2018). Functional compensation between hematopoietic stem cell clones in vivo. EMBO Rep. 19, e45702. 10.15252/embr.201745702.

45. Bramlett, C., Eerdeng, J., Jiang, D., Lee, Y., Garcia, I., Vergel-Rodriguez, M., Condie, P., Nogalska, A., and Lu, R. (2023). RNA splicing factor Rbm25 underlies heterogeneous preleukemic clonal expansion in mice. Blood 141, 2961–2972. 10.1182/blood.2023019620.

46. Jiang, D., Chowdhury, A.Y., Nogalska, A., Contreras, J., Lee, Y., Vergel-Rodriguez, M., Valenzuela, M., and Lu, R. (2024). Quantitative association between gene expression and blood cell production of individual hematopoietic stem cells in mice. Sci. Adv. 10, eadk2132. 10.1126/sciadv.adk2132.

47. Zheng, C.-X., Chen, J., Tian, J.-Y., Huang, X.-Y., Jin, Y., and Sui, B.-D. (2022). Optimized immunofluorescence staining protocol for identifying resident mesenchymal stem cells in bone using LacZ transgenic mice. STAR Protoc. 3, 101674. 10.1016/j.xpro.2022.101674.

